# Reduction of cellular toxicity with hybrid nanoparticles for mRNA delivery

**DOI:** 10.1101/2025.03.11.642711

**Authors:** Vladimir Riazanski, Lada Purvina, Luca Cavinato, Zihao Sui, Lauria Sun, Deborah J. Nelson

## Abstract

Building on the success of COVID-19 vaccine development, lipid nanoparticles (LNPs) have emerged as leading vehicles for mRNA delivery in a range of therapeutic applications. Naturally-occurring extracellular vesicles (EVs), which share similar physical properties with LNPs, present a promising alternative platform because of their relative stability and lower immunogenicity. A key challenge common to both EVs and LNPs is enabling efficient vesicle - cell interactions and establishing a polarized permeability pathway required for effective cargo transfer. Membrane recognition and intercalation are essential for the function and delivery capacity of both systems, regardless of their complexity. In this study, we leveraged recent advances to create hybrid extracellular vesicles (HEVs) by using LNPs to load mRNA into EVs. We characterized HEV formation using Förster resonance energy transfer (FRET), cryo-electron microscopy (Cryo-EM), and super-resolution microscopy, and demonstrated their ability to deliver mRNA to recipient cells. In both, *in vitro* and *in vivo* models, HEVs exhibited superior transfection efficiency compared to conventional LNPs composed of synthetic lipids, while significantly reducing LNPs cytotoxicity - a not-well-recognized limitation of synthetic lipid-based systems. These results highlight HEVs as a safer and more effective alternative for mRNA and small molecule delivery. Future therapeutic strategies could involve isolating EVs from patients, hybridizing them with synthetic lipid carriers loaded with therapeutic cargo, and reintroducing them for personalized treatment.

## Introduction

Extracellular vesicles (EVs) are membrane-bound structures with a membrane lipid content and topology identical to the cells from which they are derived. They are recognized as candidate therapeutic agents as well as drug delivery vehicles by properties including biocompatibility, low immunogenicity, negligible toxicity, immune priming, homing/targeting, cargo diversity and capacity as well as apparent longevity in circulation. Despite the absence of a mechanism enabling fusibility, they have been implicated in a diversity of roles ranging from tissue homeostasis to disease development and progression by apparent paracrine effects. It is possible that such vesicle-target cell communication occurs through EV surface proteins and their recognition or interaction with plasma membrane proteins of target cells. This proposed interaction, which is reportedly highly diverse and unique to the model under investigation, could and can result in the initiation of intracellular signaling-cascades or a change in target cell phenotype achieved through a transfer of the lumen-enclosed and/or membrane-embedded EV cargo into the cells ^1^ or simply as a function of surface receptor activation in the target cell. The literature surrounding the function of EVs in disease is immense where they have long been recognized as heterogeneous carriers for genetic information throughout the body with cell specific microRNA sorting mechanisms ^2^. Recent evidence suggests that EVs are derived from lipid raft microdomains in the plasma membrane containing proteins and nucleic acids at higher concentration than the cytoplasm of the cells from which they are derived ^3 4^. This lipid bilayer complexity, thus, suggests that EVs have the ability to deliver condensed packages of exogenous biologically active macromolecules that remain protected from enzymatic degradation while circulating in the transporting soluble milieu ^3^. They are, unlike LNPs, not immunogenic, are non-toxic, and apparently carriers capable of maintaining their own stability and of mRNA cargo. Hybrid LNP-EVs (HEVs) are theoretically capable of delivery of both soluble and membrane bound cargo, simultaneously ^5^.

Synthetic lipids including DOTAP, DOPE, MC3, and PEG lipids are used in LNPs for delivering anti-viral vaccines, similar to transfection agents for DNA transport. These LNPs use multi-layered lipids to induce fusion or membrane disruption with low toxicity, immunogenicity, and enhanced stability. Exosomes/vesicles can be modified by fusing them with synthetic liposomes to enable non-specific cellular uptake ^6 5 7 8 9 10^. Another cargo delivery strategy utilizing EVs involves transforming them into self-assembling enveloped protein nanocages (EPNs), improving cargo loading and delivery, though it relies on stochastic matching with VSV-G, with unknown but low probability ^11 12^.

In this work, we address a major challenge in developing strategies to enhance the loading and delivery of nucleic acids and proteins by extracellular vesicles (EVs) to targeted cells. We refer to “liposomes” as empty lipid bilayer vesicles and use the term “LNPs” for lipid nanoparticles encapsulating mRNA, whether composed of single or multiple lipid types. To further validate EV incorporation, we also utilized other inert nanoparticles, including fluorescent nanobeads. We systematically tracked and documented the fusion of EVs with both empty liposomes and mRNA-loaded liposomes (LNPs) as a promising approach to engineer membrane fusion. To confirm that fusion occurred specifically between EVs and LNPs, rather than as aggregates, we employed fluorescence resonance energy transfer (FRET), super-resolution microscopy, and cryo-microscopy. Although fusion events were observed at a statistically low frequency, these modified EVs, designated as hybrid EVs (HEVs), significantly enhanced gene expression levels both in vitro and in vivo.

## Results

Experiments were carried out using isolated exosomes from HEK293T and J774.1 cells grown in attached cultures. We harvested EVs from both cell lines using a standard centrifugation protocol schematized in Fig. 1A. Dynamic Light Scattering (DLS) and Nanoparticle Tracking Analysis (NTA) were used to verify concentration range and particle size in some of the experimental studies (Fig.1D). Isolated populations of EVs were further characterized in immunoblots validating the presence of the EV membrane markers, CD63, CD81, TSG101 (Fig. 1B). Immunogold cryo-images of anti-CD63 further validated the presence of the tetraspannin in Fig. 1C1. In Cryo-EM images, most exosomes displayed a characteristic bilayer membrane enclosing denser luminal cargo (Fig. 1C2).

**Figure 1.**
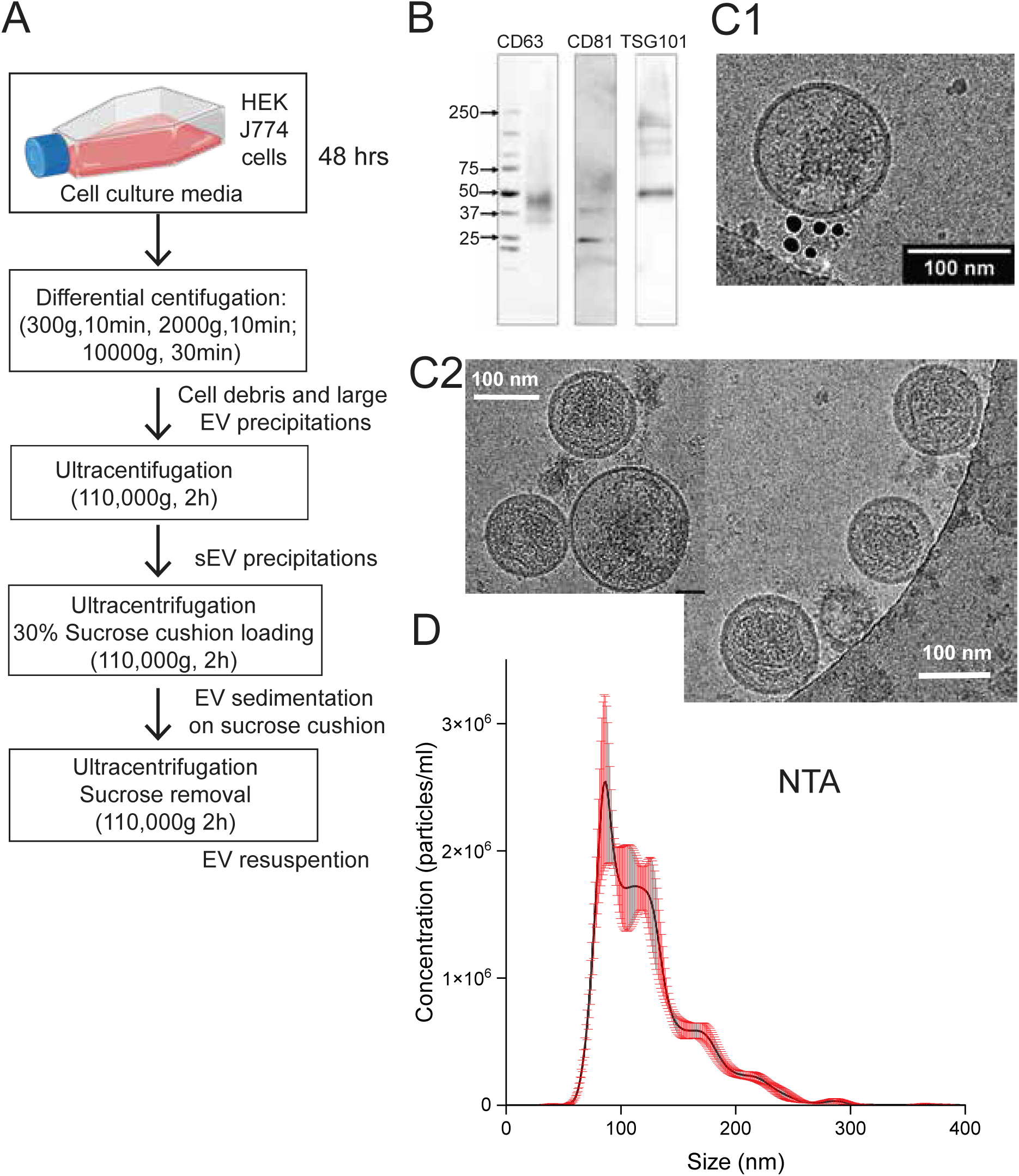
EV separation and characterization. (A) Stages of centrifugation based EV isolation technique. HEK293T and J774 cell culture lines media harvested every 48 hours undergone through various centrifugation steps. Initial differential centrifugation steps: 300g 10 min; 2000g 10 min; 10,000g 30 min were designed to remove cell debris and large EV precipitations. Following ultracentrifugation step (110,000g for 2 hours) was designed to remove small EV (sEV) precipitations, than an ultracentrifugation (110,000g for 2 hours) was performed in order to sediment EVs on 30% Sucrose cushion. That was followed by an ultracentrifugation step (110,000g for 2 hours) designed for Sucrose removal followed by EV resuspension. (B) Western blot analysis of EV markers: CD63, CD81, TSG101. (C) Cryo-EM pictures of EV samples: (C1) An EV labelled with anti-CD63 immunogold antibodies. (C2) Cryo-EM pictures demonstrating representative EV morphologies. (D) A graph showing distribution of EV sizes analyzed with Nanoparticle tracking analysis (NTA) technique.

The transfer of an EV associated transmembrane protein or luminal cargo to a cell of interest requires a fusion event. Fusion between EVs, empty liposomes, as well as complex LNPs and the plasma membrane and endosomal membranes does not happen spontaneously as the cell’s lipid bilayer is a stable structure, requiring a protein intervention to disrupt the initial bilayer configuration and facilitate mixing of the membrane lipids between the two structural elements.

### R18 as a fluorescent indicator for fusion between EVs and recipient cell membranes in the presence of PEG-1500

To gain insight into the process of EV membrane fusion with a target cell membrane, isolated EVs were loaded with a membrane fluorescent probe, Octadecyl Rhodamine B Chloride (R18). Single vesicle fusion events were detected as an increase in R18 fluorescence intensity resulting from dilution of the self-quenched probe with the cell plasma membrane. With Cell Mask dye to outline cell boundaries and EV labeling with R18, we tracked single nanoparticles during live-cell imaging. This allowed visualization of their interaction with recipient cell membranes to detect individual vesicle fusion events. Unexpectedly, we found that a simple interaction with the cellular membrane did not lead to EV fusion, even after prolonged surface contact for up to 10 minutes (Fig. 2A-C). Fusion events were observed only after the addition of the fusogenic agent PEG-1500. Fluorescence intensity peaked approximately 100 seconds after PEG addition to the recording media and declined to baseline levels over the next 10 minutes (Fig. 2B-C).

**Figure 2.**
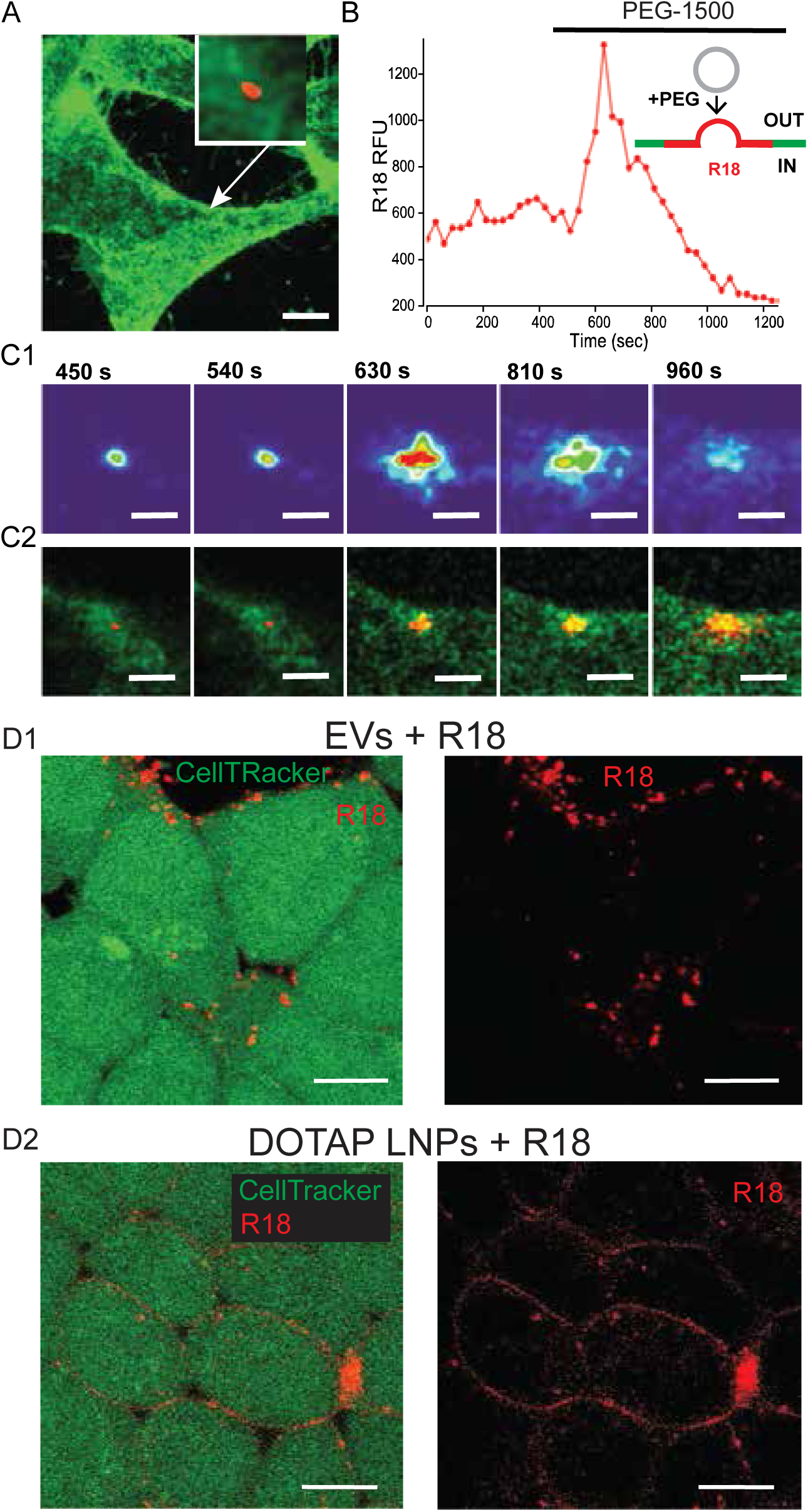
A fusogen requirement for EVs fusion with the cell membrane of the recipient cell. (**A**) An image of Octa-decyl Rhodamine B Chloride (R18) labelled EV (red) on a CellMask Green labelled HEK293T cell (green). Scale bar: 10 µm. (**B**) Addition of a fusogen polyethylene glycol (PEG-1500) initiates the fusion process of EVs to cell membrane that can be detected by dequenching related increase in the R18 fluorescence intensity followed by diffusion related decay of the signal intensity. (**C**) The time course of the EV to cell fusion process: (**C1**) – R18 mean fluorescent intensity change in heat map lookup table; (**C2**) – spreading of R18 (red) label on the surface of CellMask Green labelled HEK cell (green) over time. Scale bar: 2 µm. (**D**) The comparative difference of R18 membrane label spread in two nanoparticle preparations: **D1** – R18 labelled EVs (red) after addition to HEK cells labelled with CellTracker Green (green) , **D2** - R18 labelled DOTAP LNPs after addition to HEK cells labelled with CellTracker Green (green). Scale bar 5 µm.

Notably, no fusion events were detected for over an hour before the addition of PEG, confirming that a fusogen is required to initiate and facilitate fusion of EVs with cell plasma membrane. The decline in fluorescence intensity corresponded to the diffusion of the dye within the bilayer of the recipient cell membrane. These data provided a time course for the vesicle-cell fusion process. Having validated the PEG-facilitated fusion timeline, we compared vesicle interactions with cell membranes between R18-labeled EVs and DOTAP liposomes at higher resolution in the absence of PEG. As observed in previous experiments, prolonged incubation (10–15 minutes) of R18-labeled EVs with HEK293T cells did not result in plasma membrane labeling. Instead, R18 fluorescence appeared as small puncta (Fig. 2D1) either on the cell surface or intracellularly presumably due to endosomal uptake. In contrast, the addition of R18-labeled DOTAP liposomes led to plasma membrane labeling that outlined cell surfaces, indicating that the fluorescent dye spread across the cell surface as a result of vesicle fusion (Fig. 2D2).

These experiments demonstrated that, in the absence of membrane protein fusogens, EVs are unable to fuse with cellular membranes. This observation suggested to us that incorporating synthetic lipids, similar to those used in LNPs, into the EV membrane may be necessary to facilitate cargo delivery to recipient cells.

### Liposomes, LNPs and HEV preparation

We used synthetic lipids to prepare both liposomes and LNPs. Liposomes (Lip) were prepared without cargo using the extrusion method and were used primarily for toxicity and FRET studies to examine fusion with EVs. The synthetic lipids used were chosen based on descriptions of LNP formulations in delivery systems for generic drugs, mRNA, and small interfering RNA, similar to those used in Covid-19 vaccines ^13^. A review of the LNP therapeutic field from Cullis *et al.* further informed our synthetic lipid choices and our understanding of the LNP delivery systems ^14, 15^. Our studies encompassed the use of the cationic lipid DOTAP. Currently, several techniques are available for forming hybrid vesicles by fusing EVs and liposomes. In the literature there are multiple techniques for forming hybrid vesicles including simple mixing ^16^ and electrostatic interaction of oppositely charged particles (for review ^17^).

### Hybrid vesicle preparation

Nanoparticle electrostatic interactions significantly influence the behavior of charged particles, impacting the stability and aggregation of various nanoparticle formulations. It has been proposed that simple mixing of EVs and synthetic vectors can lead to their fusion due to the attraction between different charges. To investigate this, we measured the zeta potential of cationic DOTAP liposomes and J774.1-derived EVs before and after the extrusion process, which results in hybrid extracellular vesicles (HEVs) (Fig. 3B).

**Figure 3.**
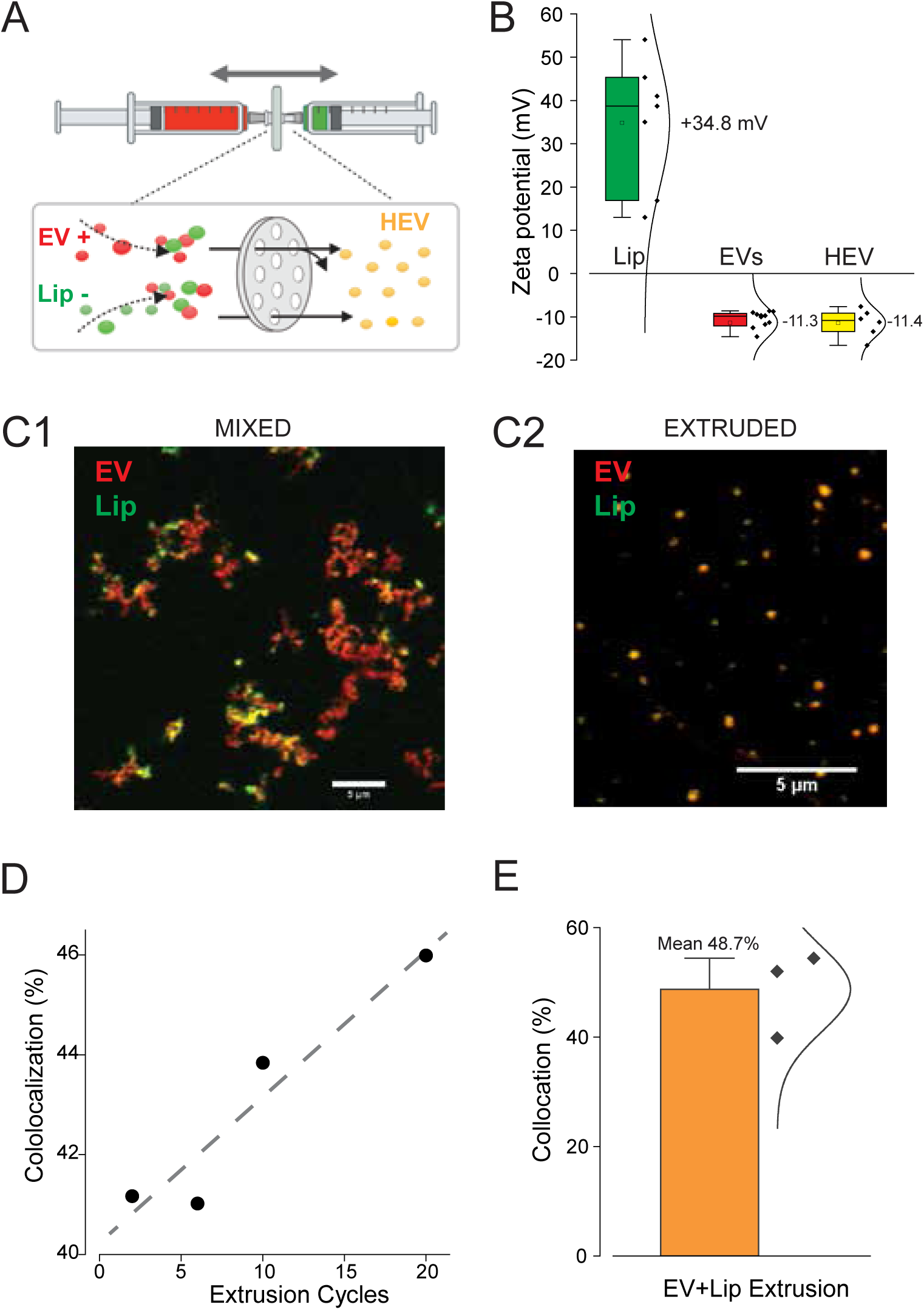
HEV formation using extrusion methodology. (**A**) Schematic representation of the extrusion method for obtaining hybrid extracellular vesicles (HEV, yellow) by multiple passing through a filter Bodipy-TR (red) labelled EVs and NBD-PE fluorescent (green) lipids labelled Lip. (**B**) Zeta potential measurements summary graph of: freshly made DOTAP liposomes (Mean ζ=+ 34.8mV, n=7 samples), J774 EVs (mean ζ=-11.3 mV, n=12 samples) and HEVs (mean ζ=-11.4 mV, n=6 samples). (**C**) Confocal microscopy images of mixed EVs (red) and Lip (green) before (**C1**) and after 20 cycles of passing through extrusion filter. Scale bar: 5μm. (**D-E**) Colocalization analysis, particle size distribution, and polydispersity index after the formation of hybrid particles. Quantification was determine using Image-J and Origin software.

DOTAP liposomes and EVs exhibited significantly opposite charge, with the liposomes having a positive and more variable zeta potential (average +34.8 mV), whereas the EVs had a negative zeta potential (average -11.3 mV). The zeta potential distribution was considerably broader for the DOTAP liposome preparation compared to EVs and HEVs, indicating lower stability of the lipid nanovesicle formulations, likely due to the Ostwald ripening process ^18^. After 20 extrusion cycles, the resulting HEV particles exhibited a zeta potential very similar to that of EVs, averaging -11.4 mV. Despite mixing liposomes and EVs in a 1:1 ratio, the resulting charge closely resembled that of EVs alone, potentially due to the higher mass of EVs relative to liposomes.

To assess hybrid vesicle formation, we performed a colocalization analysis using different fluorophores to label the nanoparticles. EVs were stained with Bodipy-TR (red), while liposomes were labeled with NBD-PE (green) fluorescent lipids (Fig. 3A, C). Mixing the positively charged liposomes with the negatively charged EVs led to particle aggregation due to electrostatic interactions, which was visualized through confocal imaging (Fig. 3C1) with significant color colocalization. Following the extrusion of the liposome and EV mixture in a 1:1 proportion, the percentage of colocalization increased linearly with the number of extrusion cycles (Fig. 3D), reaching 48.7% after 20 cycles (Fig. 3E). These results suggest that electrostatic attraction is sufficient to cause particle aggregation between EVs and liposomes, but particle fusion requires the application of shear force through the extrusion process.

### FRET validation of HEV production

Despite the promising colocalization results, the formation of hybrid nanoparticles smaller than the light diffraction limit (>200 nm) cannot be reliably distinguished from particle aggregation. To address this issue, we employed Förster Resonance Energy Transfer (FRET), a technique previously used to evaluate hybrid vesicle formation ^6, 10 5^. We found that the FRET pair consisting of NBD-conjugated lipids in liposomes (donor) and Bodipy-TR ceramide labeling in EVs (acceptor) forms an efficient FRET system for detection of HEV formation (Fig. 4A). The technique can be used to monitor either a decrease in the donor signal (donor quenching - DQ) or an increase in the FRET signal following extrusion-induced hybridization (Fig. 4B2). A significant reduction in the NBD donor signal, accompanied by an increase in FRET efficiency, was observed after the formation of HEVs (Fig. 4C1-C2).

**Figure 4.**
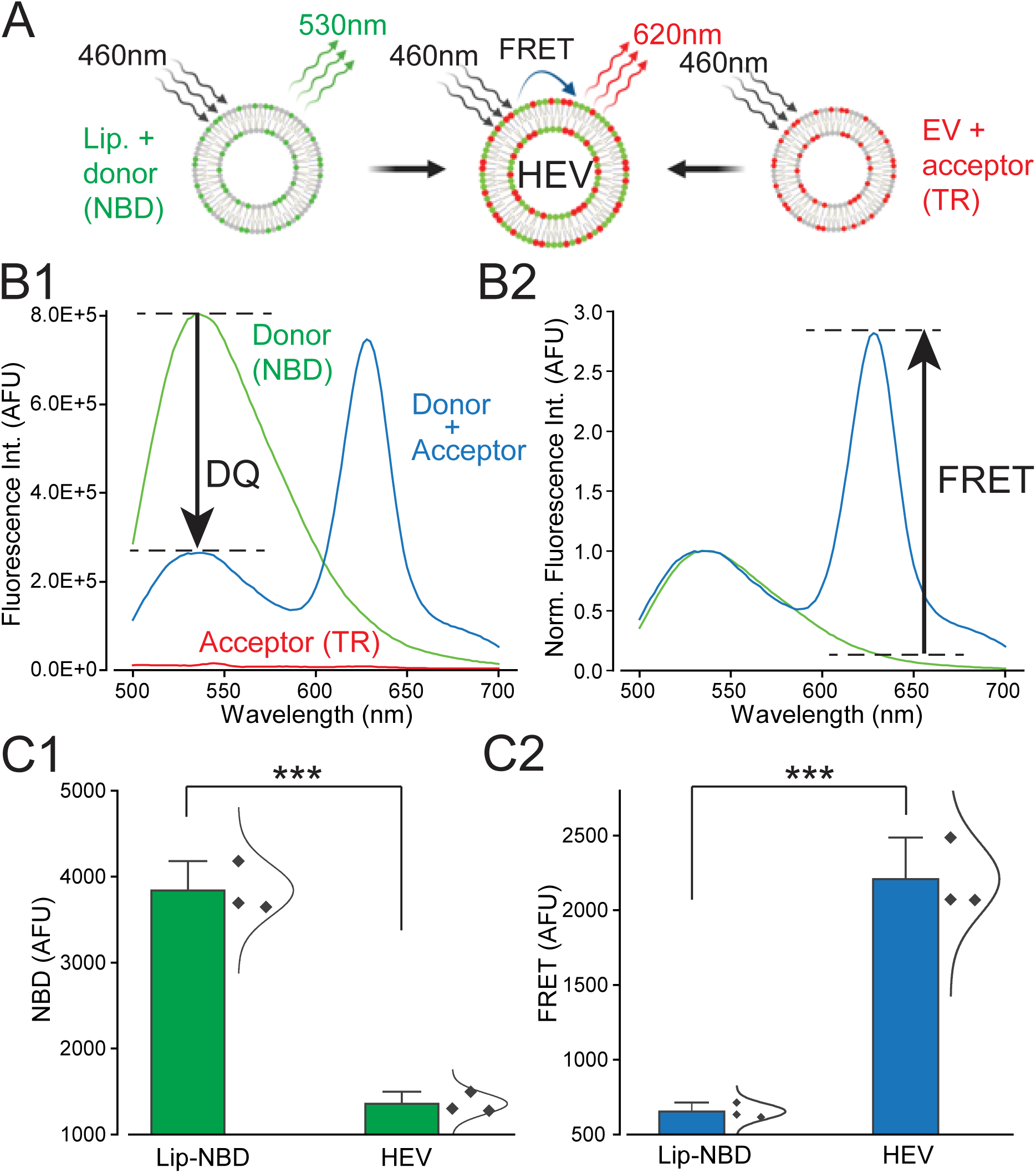
Validation of HEV production with FRET. (**A**) Schematic representation of FRET signal transfer during the fusion of DOTAP liposomes labeled with FRET donor NBD and EVs labeled with FRET acceptor Bodipy-Texas Red (TR). Dye stock solutions were 1 mM with a dilution of 1:100 for experimental use. (**B1**) Fluorescence spectral emission scans during excitation at 460 nm: donor alone (NBD, green line); acceptor alone (TR, red line); and donor plus acceptor after formation of HEVs (blue line). An arrow indicates the decrease in donor peak emission as a result of donor quenching (DQ) during FRET. (**B2**) The spectral scans normalized to the donor peak reveal a sensitized acceptor emission component due to FRET (black arrow). (**C1**) Analysis of the decrease in donor signal at Ex 460/Em 530 nm (Lip-NBD n=3; HEV n=3). (**C2**) Increase in FRET signal intensity following EV-liposome fusion, as determined by plate reader at Ex. 460nm / Em. 620nm (Lip-NBD n=3; HEV n=3).

### EV/Lip/HEV visualization by cryo-electron microscopy

Although delivery of nanoparticle cargo via endosomal escape has been reported to be dependent on proteins and pH which can be modulated by changing the lipid structure in the endosome or at the plasma membrane of target cells, their identification remains unknown ^19 20^. According to published data from Bonsergent et al. ^19^, spontaneous EV endosomal escape is somewhere less than 1% per hour and less than 30% of available nanoparticles are taken up into acceptor cells. Thus, optimization of cargo delivery by engineered HEVs is essential to ensure a functional cellular response. If the fusion between membranes by either pH changes or lipid interactions is to be understood, examination at the EM level is essential. Our visualization of HEV structure can be seen in the cryo-EM examples of Fig. 5. Images of a DOTAP liposome, an isolated EV, the fusion process, and final hybrid are displayed in Fig. 5A. Liposomes were distinguished by their uniform, characteristic round shape and empty lumen. In contrast, EVs exhibited a less spherical surface and, in most cases, contained visibly dense luminal cargo. This morphological difference allowed us to differentiate between the two types of vesicles and to detect their interactions, which led to the formation of a putative fusion pore (Fig. 5D) between the vesicles, ultimately resulting in full fusion and the formation of hybrid vesicles. In the resulting HEVs, the fused liposome appeared as a bleb on the EV surface (Fig. 5E). Similar vesicle fusion events have been reported elsewhere ^20^ .

**Figure 5.**
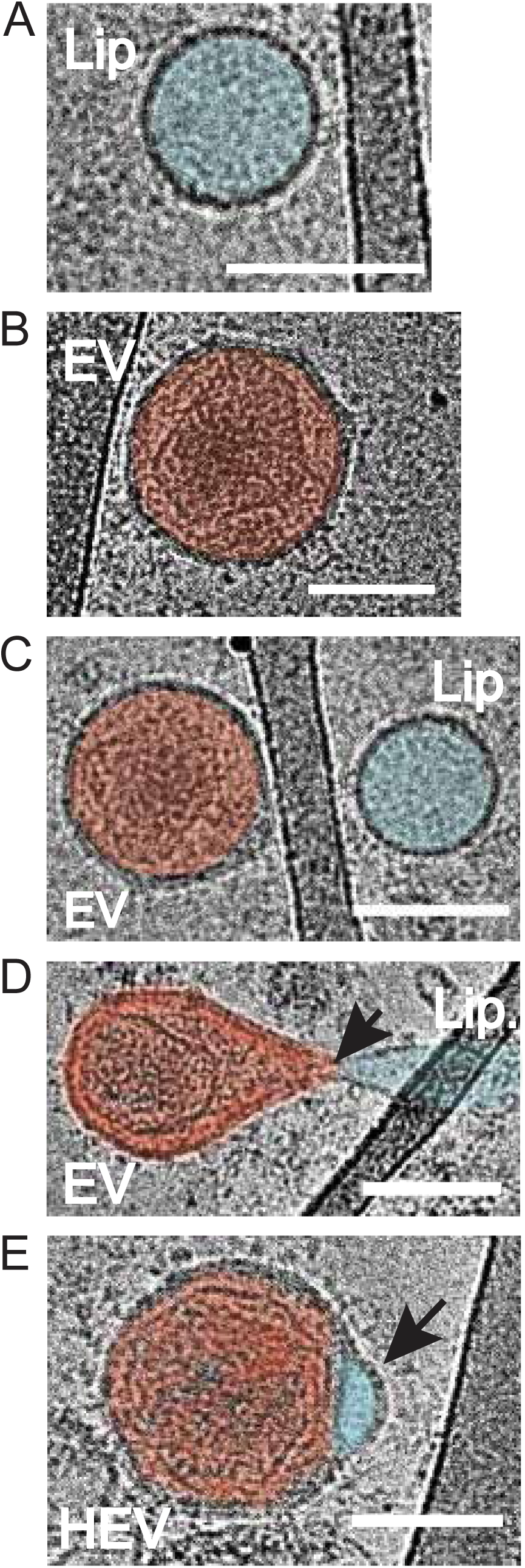
High-resolution Cryo-EM imaging of HEV formation. (**A**) Cryo-EM images of DOTAP liposomes (Lip, cyan pseudo coloring) and EVs (EV, red pseudo coloring) before HEV formation (**A-C**) and during fusion (**D-E**). The black arrowhead indicates the fusion pore. (**E**) The resulting hybrids after fusion (HEV) have characteristic ear-like bumps indicated with a black arrowhead. Scale bars are 100 nm.

Additionally, super-resolution stimulated emission depletion (STED) microscopy was used to analyze the colocalization of LNP loaded with 100 nm fluorescent FluoSpheres (FS) and EVs labeled with Abberior STAR RED™ membrane dye (Fig. 4B). The resulting HEV particles displayed signals from both the EVs and the fluorescent beads, indicating successful fusion and concomitant bead incorporation forming a successful HEV construct. Similar to the Cryo-EM images of HEV formation, the STED images showed that the fluorescent beads were localized off-center within the HEVs, suggesting potential spatial limitations in the lumen of the fused vesicles (see inclusions in the EVs of the Cryo-EM images in Fig. 5). Gaussian fittings of the full width at half maximum (FWHM) measurements from deconvoluted images revealed sub-diffraction-limited sizes for FS, EVs, and the resulting HEVs (Fig. 6A3).

**Figure 6.**
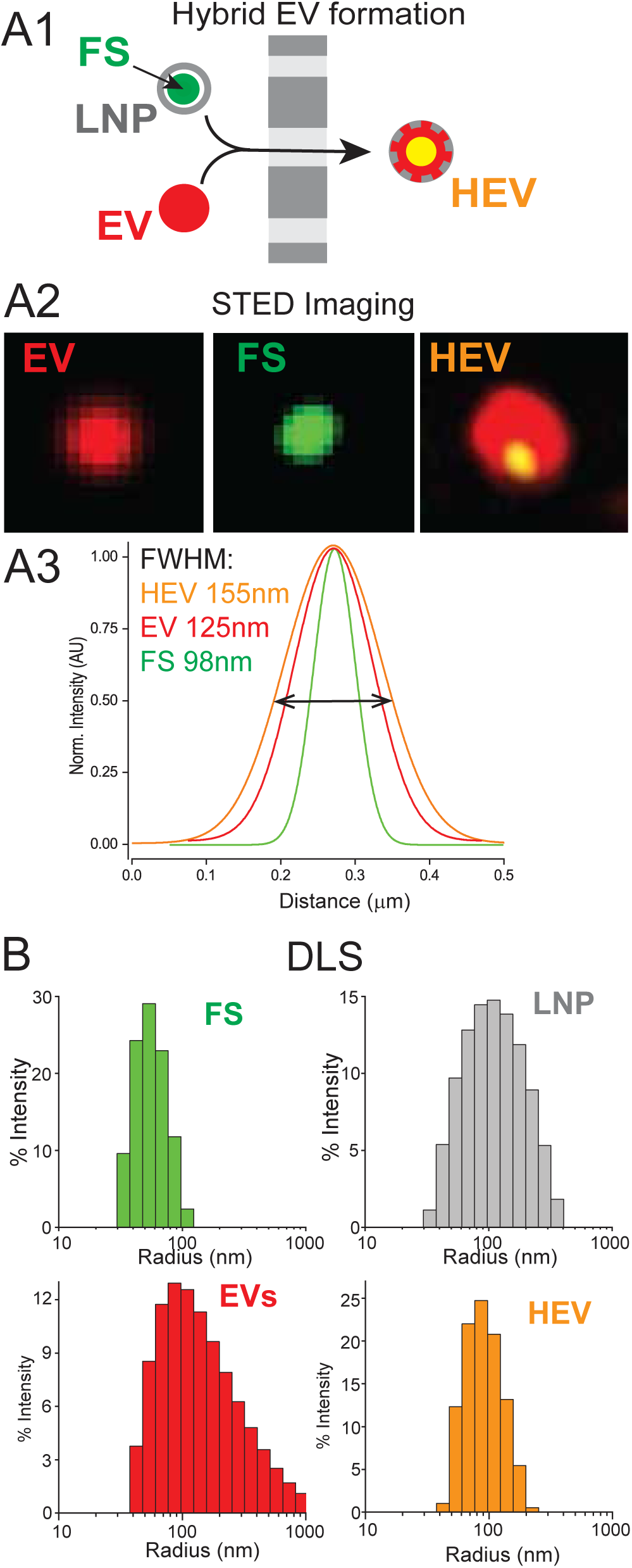
(**A1**) A schematic representation of HEV preparation with extrusion of fluorescently labeled EVs (EV, red) and DOTAP LNP with 100 nm encapsulated FluoSpheres (FS, green). (**A2**) Super-resolution STED imaging of fluorescently labeled EV (EV, red), FS (green) and HEVs with encapsulated FS (red and yellow) after deconvolution. (**A3**) The graph illustrates Gaussian fittings of the three nanoparticles’ fluorescence intensities for size estimation by measuring the full width at half maximum (FWHM). (**B**) Dynamic light scattering (DLS) measurement of the nanoparticles’ size distributions in bulk solution: FluoSpheres (FS, green); DOTAP LNP (LNP, grey); EV (EV, red); HEVs (HEV, orange).

Size distribution measurements and analyses provided further insights into the physical characteristics of the hybrid particles. Dynamic light scattering (DLS) measurements of nanoparticle size distribution indicated that HEVs exhibited narrower size distributions compared to EVs or liposomes (Fig. 6B), suggesting that extrusion primarily determines the size of the hybrid vesicles based on the pore size of the extruder, rather than the original vesicle sizes.

### Comparative Lip, LNP and HEV cellular toxicity *in vitro*

HEVs hold promise in overcoming rapid clearance of mRNA as well as siRNA delivery vehicles *in vivo*. Their promise also includes enhanced stability and decreased toxicity over that seen with that seen with multi-component LNPs alone. Initially, comparative toxicity of the nanoparticles in our study was assessed *in vitro* using cultured HEK cells. For these studies we used the CyQUANT^TM^ XTT Cell Viability assay. Cells were seeded a day before the assay on 96 multi-well plates then exposed for 3 hours to an increasing number of either EVs, liposomes, LNPs or HEVs, fused with DOTAP liposomes. HEVs were derived by extruding an admixture of 50% purified EVs and 50% of DOTAP liposomes. Complex LNPs were composed of the complex lipid formulation (Avanti Lipid Blend 2: 38.5%, Sitosterol, 49% DOTAP 11 mol% DSPC, 1.5 mol% DMG-PEG 2000) available commercially. Cell viability was reported as percentage of survival of treated-cells compared to untreated ones (Fig. 7B). Complex LNPs and liposomes significantly reduced cell viability in a concentration-dependent manner (e.g., at 10⁷ and 10⁸ particles), whereas, cells treated with EVs or HEVs exhibited exceptionally low toxicity, even at the highest concentrations tested. These results were also confirmed, but not quantified, using a live-cell viability assay, imaging propidium iodide (PI) of the nanoparticles (see microscopy images in Fig. 7A).

**Figure 7.**
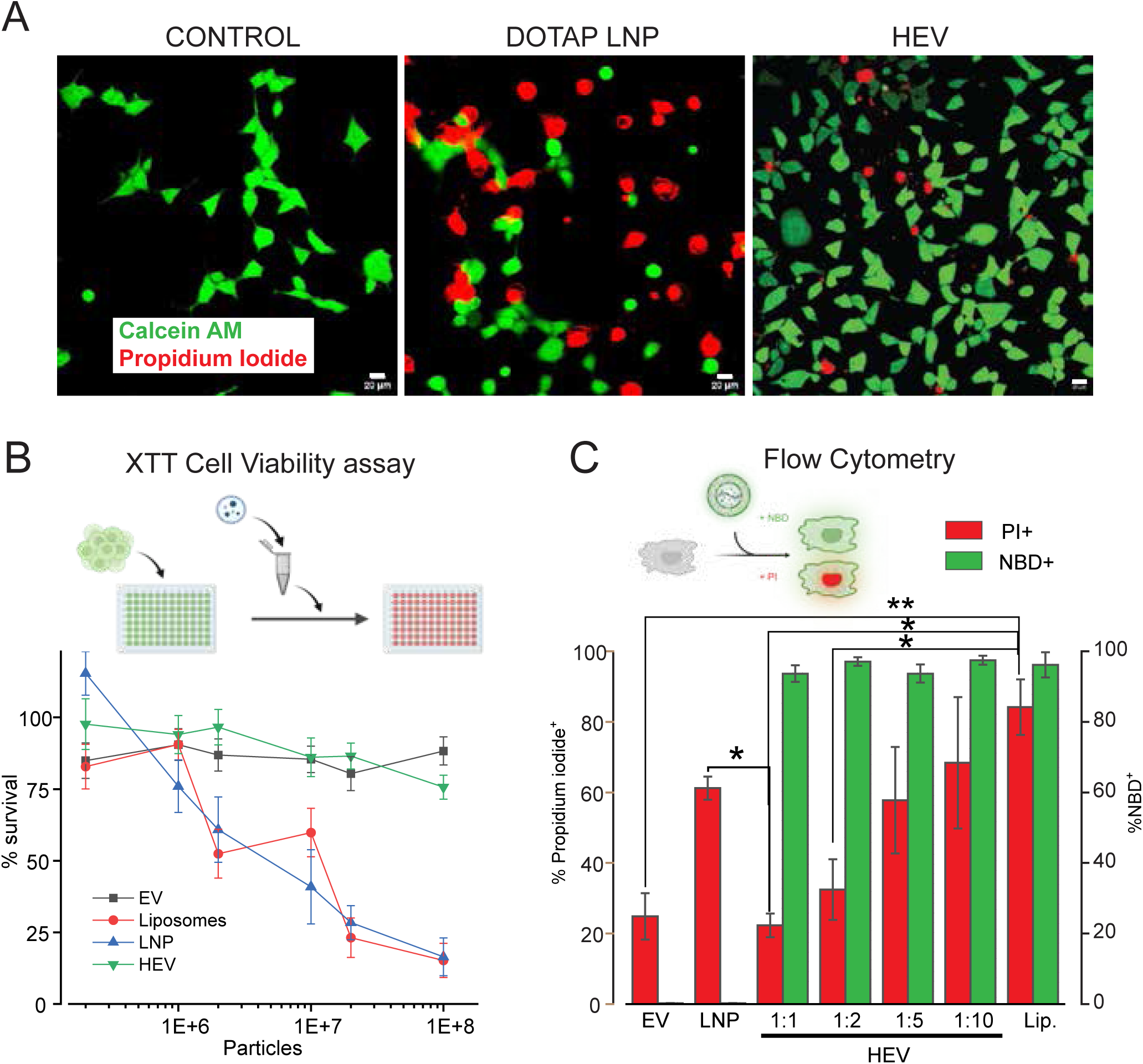
In vitro viability and delivery assays to assess HEV toxicity and efficacy. (**A**) Representative fluorescence microscopy images of HEK293T cells stained with Calcein AM (green, viable cells) and Propidium Iodide (red, dead cells) 1 hour after treatment with artificial cerebrospinal fluid (Control), DOTAP LNPs, or HEVs. Scale bar 20 (**B**) CyQUANT XTT Cell Viability Assay shows HEK293T survival when exposed to increasing numbers of the indicated particles: EV (black), liposomes (red), LNP (blue), and HEV (green). The percentage of survival was calculated relative to untreated cells. (**C**) Flow cytometry analysis of HEK cell viability and liposome delivery efficacy. Toxicity was evaluated as the percentage of propidium iodide-positive cells (red bars), and efficacy was assessed in the bulk live population as the percentage of NBD-positive cells (green bars). Statistics: Two-way ANOVA (A) or One-way ANOVA (B) with Bonferroni multiple comparisons. **, p<0.01; *, p<0.05. n = 4 for each experimental condition.

Similar cell viability results were obtained using flow cytometry of HEK cell cultures treated with nanoparticles labeled with fluorescent phosphoethanolamine (NBD-PE). After a 3-hour incubation, cells were dissociated and stained with 1 µM PI to assess cell death within the population of NBD-labeled cells. The results confirmed higher toxicity of complex LNPs and DOTAP liposomes as compared to EVs and HEVs. Notably, HEV toxicity was dependent on the proportion of DOTAP lipids used in hybrid formation, with toxicity increasing in a ratio-dependent manner (Fig. 7C).

### Comparative Lip, LNP and HEV cellular toxicity *in vivo*

Multiple therapeutic approaches are being developed to deliver mRNA to lung cells for the treatment of diseases such as cystic fibrosis and COPD ^21 22 23^. This underscores the need to evaluate the cellular toxicity of mRNA delivery nanoparticles administered via the intratracheal route.

Nanoparticles at a concentration of 1×10^8^ /mL (the dose that showed the greatest difference in the *in vitro* XTT cell viability assay) were administered together with 100 µM PI to evaluate cell death during a one-hour exposure to nanoparticle-containing solutions in a mouse *in vivo* assay. Dead and live cell nuclei were distinguished in exposed lung slices via NucBlue staining and quantified (Fig. 8A-B). The majority of cell damage in the *in vivo* preparation was observed along the bronchiolar walls, consistent with trends noted in the *in vitro* assay. Cell death was reported as the percentage of propidium iodide-positive nuclei relative to the total number of NucBlue-stained nuclei in the regions adjacent to the bronchiolar space. After one hour of exposure, DOTAP liposomes and complex LNPs significantly increased lung cell death, whereas EV- and HEV-treated lung slices exhibited exceptionally low toxicity, comparable to the control injection of ACSF (Artificial Cerebrospinal Fluid) alone (Fig. 8C). The lower cell death rate compared to the *in vitro* assay may be explained by the shorter exposure time and reduced bronchial tissue permeability, likely due to mucosal surface coatings ^24^.

**Figure 8.**
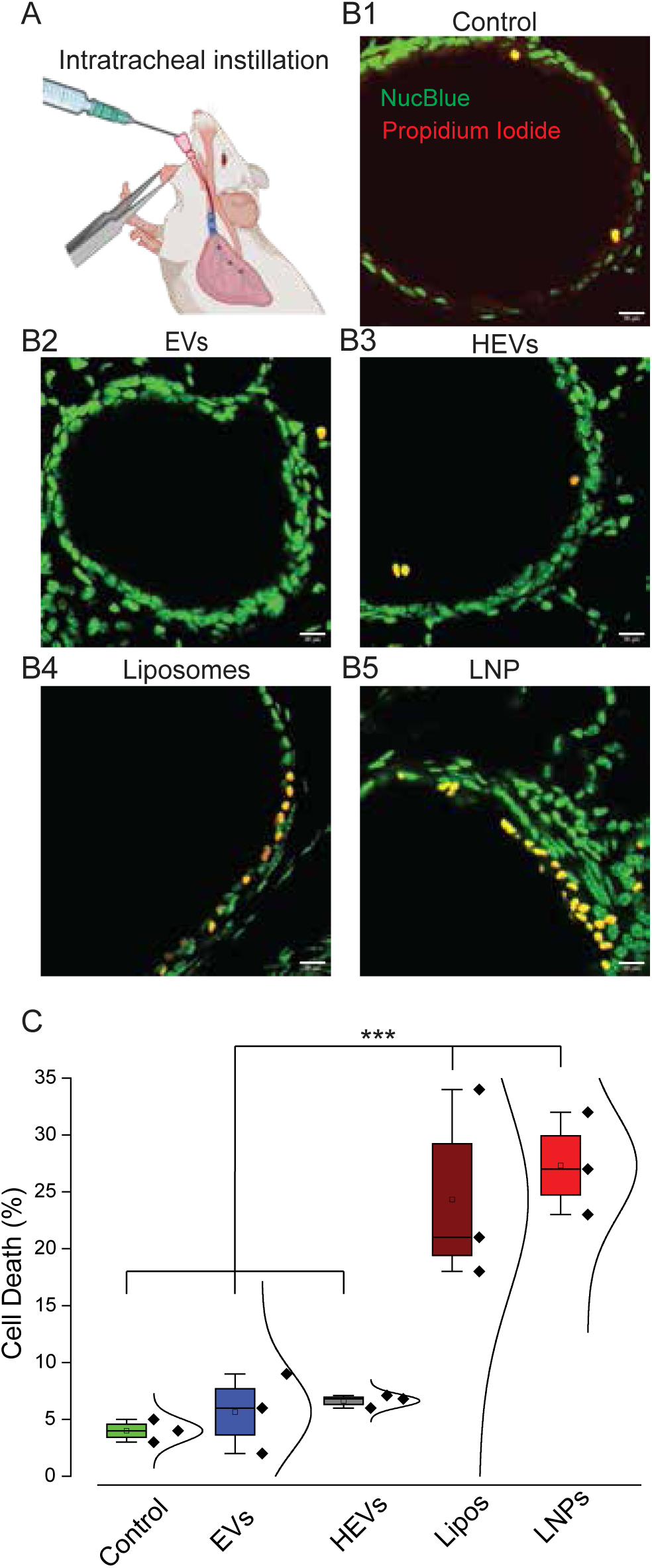
LIVE/DEAD cell viability assay on acute lung slices isolated from mice after mouse intratracheal instillations. (**A**) A cartoon depiction of mouse intratracheal instillation. (**B**) Representative acutely cut lung sections containing part of a bronchiole stained with NucBlue (green) and Propidium Iodide (red) after injections of: (**B1**)-ACSF (Control), (**B2**) - 1x108 /ml EVs, (**B3**) - 1x108 /ml HEVs, (**B4**) - 1x108 /ml DOTAP Liposomes, (**B5**) - 1x108 /ml Moderna LNPs. Scale bar 20µm. (**C**). Quantification and comparative analysis of the Cell death percentage in lung injected with: ACSF (Control, green box, n=3 mice); EVs (blue box, n=3 mice); HEVs (grey box, n=3 mice); DOTAP liposomes (Lipos, maroon box, n=3 mice) and LNPs (LNPs, red box, n=3 mice).

### Comparative mRNA transfection efficiency *in vivo*

The cell cytotoxicity assays have revealed that DOTAP liposomes as well as LNPs made of complex lipids both have comparable high toxicity relative to EVs. Hybridization of liposomes with EVs diminishes the cellular toxicity of nanoparticles made with synthetic lipids, making HEVs a less toxic clinical alternative to currently used LNPs. The comparative efficacy of HEVs and their ability to deliver its mRNA cargo to recipient cells is of equal importance. To address this comparison, we used two fluorescent proteins (FPs): EGFP and mCherry. The mRNAs encoding the fluorescent proteins were encapsulated into DOTAP liposomes and Avanti Lipid Blend 2 LNP mixtures using a T-type jet mixer ^22^ or extruder. Resulting LNPs loaded with mCherry or EGFP mRNA were used to make HEVs using the filter extrusion process or in the case of complex LNPs with the NanoAssemblr™ Ignite™ . The LNPs and HEVs were added to HEK cell cultures in comparable concentrations (1x10^9^-1x10^10^ particles/ml), incubated for 3 hours and imaged after 24 hours with confocal laser microscopy for evaluation of the fluorescent proteins expression in viable cells labelled with Calcein Green AM - live cell markers (Fig. 9). Statistical comparison of transfection efficiency between LNP-FPs mRNA and HEVs-FPs RNA showed significantly higher transfection efficiency of HEVs (Fig. 9B). These results show that while HEVs have significantly lower cellular toxicity they have better mRNA transfection efficiency relative to LNPs.

**Figure 9.**
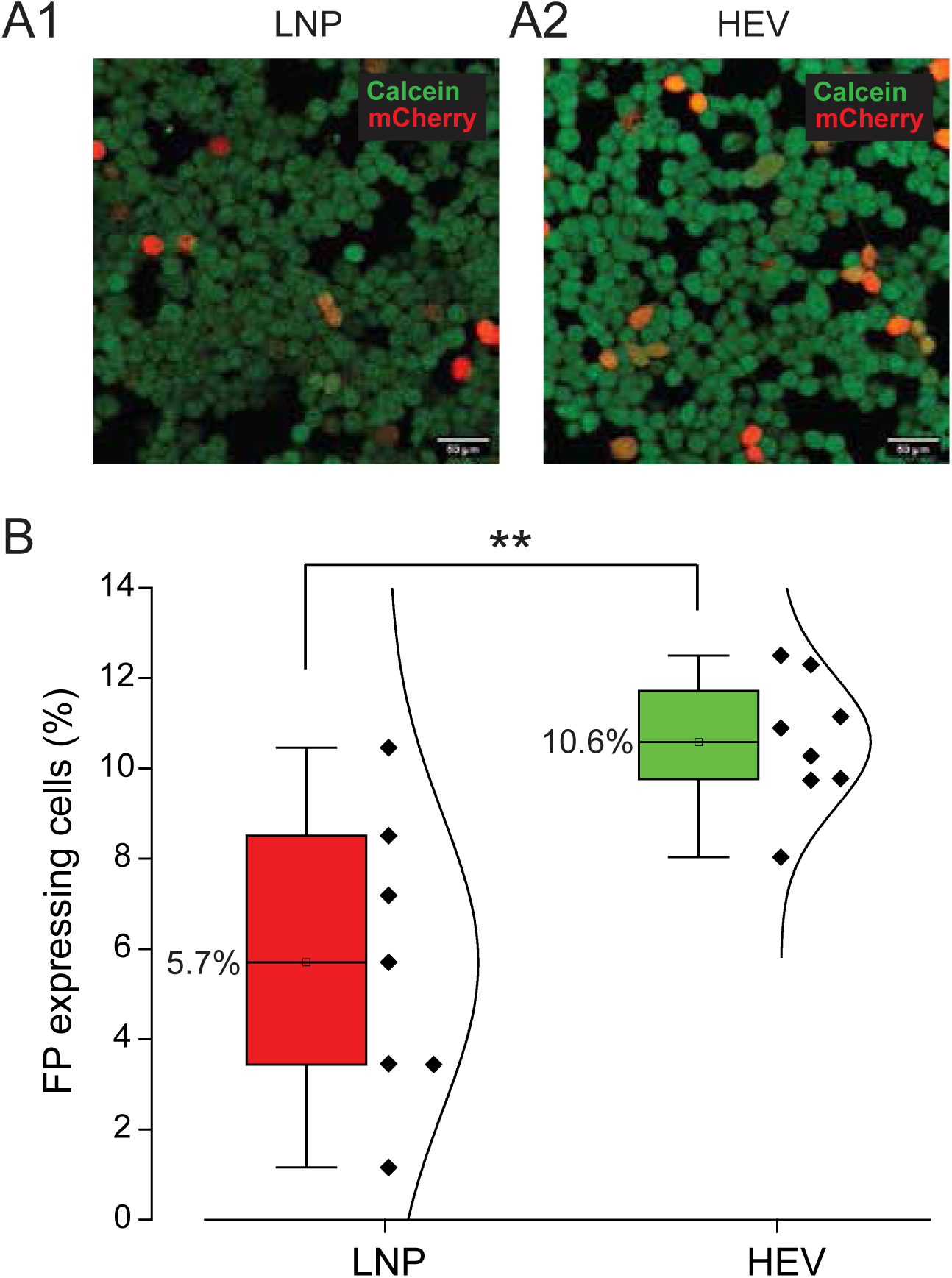
Transfection efficiency of HEK293T cells with LNP- or HEV-delivered fluorescent protein mRNA. (**A1**) Representative fluorescence microscopy image of HEK293T cells labeled with Calcein Green AM (green) 24 hours after transfection with LNP-mCherry mRNA. (**A2**) Representative fluorescence microscopy image of HEK293T cells labeled with Calcein Green AM (green) 24 hours after transfection with HEV-mCherry mRNA. Scale bar: 50 µm. (**B**) Quantification of in vitro transfection efficiency of HEK293T cells with LNPs or HEVs carrying EGFP or mCherry mRNA (LNP: mean 5.7%, n = 7; HEV: mean 10.6%, n = 8; error bars represent SE). Statistical significance was determined using one-way ANOVA (p < 0.01).

## Discussion

Hybrid extracellular vesicle-liposome nanoparticles (HEVs) represent a novel class of delivery vehicles that merge the biological advantages of natural extracellular vesicles (EVs) with the customizable design and loading capacity of synthetic lipid nanoparticles (LNPs). In this study, we developed HEVs through an extrusion approach and demonstrated their superior delivery of nucleic acids compared to parental EVs and traditional liposomes and LNPs. The resulting hybrids retained key EV features, including low immunogenicity and inherent biocompatibility, while displaying enhanced cargo encapsulation, stability, and delivery efficiency, thus addressing key challenges in gene delivery platforms ^5 8^. Our study addresses the limiting step in EV biology compromising the quantity, content, and targeting of cargo in delivery through EVs, namely, the absence of a mechanism for fusion between EVs and target cell membranes. Delivery of EV content and EV membrane proteins is dependent upon membrane fusion either at the plasma membrane or endosomal internal sites depending upon entry pathway, something that has not been reliably defined or observed at more than negligible probability to date. Taking advantage of knowledge gained in the construction and use of complex LNPs in mRNA vaccine delivery, we have investigated, formulated, and characterized hybrid lip-EVs which can be isolated from patient sources and post isolation hybridized with synthetic lipid vehicles filled with payload of choice before re-introduction to the patient. We have compared hybridization with multicomponent LNPs used in the mRNA vaccines with liposomes composed of a single lipid with its effect on cellular target fusion. It was our hypothesis that the membrane of the bio-normal EVs would provide the non-synthetic lipid components used to confer vaccine-LNP stability and extended lifetime in the circulation, thereby, reducing toxicity and enhancement of cargo delivery following membrane to membrane fusion. Given the recent plethora of studies of extracellular vesicles delivering epigenome editing systems, we assumed that the hybridization process would also facilitate fusion. To our surprise, this resulted in an extremely modest increase in delivery efficiency.

The improved functionality of HEVs may stem from their unique membrane composition, which integrates the fusogenic and targeting properties of EV membranes with the structural versatility of synthetic lipids. Prior studies have shown that lipid rafts and tetraspanin-enriched microdomains in EV membranes play critical roles in cargo sorting and membrane fusion ^4^ , and although we have not presented data validating this observation in our study, these components may similarly contribute to the efficient endosomal escape and cytosolic release observed in our hybrids. Additionally, by preserving EV-associated surface proteins, such as CD9, CD63, and CD81, HEVs may leverage receptor-mediated uptake pathways, improving their specificity and internalization by target cells ^1 3^. Importantly, the hybridization process appears to confer increased colloidal stability and protection under physiological conditions, likely extending circulation time and reducing premature clearance ^5^. These features are consistent with previous reports indicating that membrane fusion strategies can improve pharmacokinetic profiles and biodistribution compared to conventional nanoparticles ^1^. Moreover, the modular nature of HEVs may allow for surface engineering to further enhance tissue-specific targeting, a strategy that has been successfully applied in engineered EV systems ^8 5^.

In our comparison between EVs, HEVs and LNPs the most significant difference was the levels of toxicity between the three vesicle types as a function of synthetic lipid composition. Flow-cytometry studies using liposomes and LNP-exposed cells showed high rate of PI incorporation, a reporter of cell death, as compared to either HEVs or EVs. HEV toxicity was a function of the DOTAP-NBD liposome to EV ratio in the extrusion process. Surprisingly, the efficacy of cellular fusion or uptake measured as the percentage of NBD-positive cells was independent of the percentage of DOTAP incorporated into HEVs. This suggested to us that the NBD positive signal indicated HEV attachment but not necessarily fusion with the HEVs putatively hybridized with higher concentrations of DOTAP-NBD.

### Comparative LNP and HEV cellular toxicity *in vivo*

*In vitro* toxicity assessments can reveal direct cellular damage but serve primarily as an initial indicator of the dose-dependent toxicity of drug carriers. However, in one of the major target organs for LNP-based drug delivery - the lungs - the response to nanoparticle exposure can be far more complex. These reactions include, but are not limited to, activation of immune and inflammatory pathways (including complement activation and alveolar macrophage engagement), interference with surfactant function, limited clearance and accumulation, and oxidative stress. All of these factors must be accounted for in order to accurately evaluate new treatments that employ novel drug delivery systems. Therefore, we decided to assess the comparative cellular toxicity of LNPs, liposomes, EVs, and HEVs following direct lung instillation.

When formulating LNPs using synthetic lipids - especially for drug or gene delivery - it’s crucial to evaluate their potential cytotoxic effects and the mechanisms that might trigger cell death. DOTAP (1,2-dioleoyl-3-trimethylammonium-propane) is a cationic lipid commonly used to formulate lipid nanoparticles (LNPs) or liposomes for gene delivery. While it can be effective in packaging and delivering nucleic acids, DOTAP-containing formulations are known to exhibit varying degrees of toxicity, particularly in the lungs. DOTAP-containing LNPs can exhibit toxic effects in the lungs primarily due to their cationic charge, which promotes membrane disruption, inflammatory responses, and oxidative stress. Understanding these mechanisms and optimizing formulations are critical for balancing therapeutic efficacy with safety, especially for pulmonary drug and gene delivery applications.

Our study presents a compelling approach to develop and characterize hybrid extracellular vesicles as novel delivery vehicles for mRNA and protein therapeutics. The approach by hybridizing EVs with liposomes to overcome limitations of current delivery systems. Our study demonstrates that the hybrid system provides for enhanced gene delivery minimizing toxicity which would contribute to enhanced therapeutic outcomes. While our findings reinforce the potential of HEVs as effective carriers for nucleic acid therapeutics, several limitations warrant further investigation. The precise mechanisms by which hybridization enhances delivery efficacy remain incompletely understood. Specifically, dissecting the contributions of membrane lipid composition, protein cargo, and curvature-driven sorting during hybrid formation may provide deeper insights into optimizing HEV design ^1^. Future *in vivo* studies are essential to assess the biodistribution, therapeutic efficacy, and immunogenicity of HEVs in clinically relevant contexts ^2 8^. In conclusion, our study adds to the growing body of evidence supporting hybrid EV-based delivery platforms as promising tools in gene therapy and nanomedicine. By uniting the natural targeting and immune-evasive properties of EVs with the tunable features of LNPs, HEVs offer a compelling strategy to overcome critical barriers in the delivery of genetic medicines, ultimately advancing the development of next-generation therapeutics ^5^.

## Materials and Methods

### Animals

The Animal Care and Use Committee at the University of Chicago approved all the procedures outlined here. All animals were housed in a specific, pathogen-free, biohazard level 2 facility, maintained by The University of Chicago Animal Resources Center (Chicago, IL). Animal genotyping was performed by Transnetyx, Inc., (Cordova, TN).

### Mammalian Cell Culture

HEK293T and J774.1 cells were obtained from the American Type Culture Collection (ATCC). All cell cultures were maintained in CO_2_ incubators at 37°C, 5% CO_2_, and 100% humidity in high glucose containing GenClone^TM^ DMEM cell cultures medium (Genesee Scientific, El Cajon, CA USA) supplemented with 10% exosomes depleted FBS (Gibco, USA) and 1% Penicillin-Streptomycin ( Gibco, Thermo Fisher Scientific, Waltham, MA USA). For EV isolation cells were grown either in standard T75 cell culture flasks or in CELLine Bioreactors (Argos Techologies Inc, Vernon Hills, IL USA).

### EV isolation and characterization

EVs from the cell culture media were isolated using two different methods: 1) with multistage ultracentrifugation and 2) using asymmetric flow field-flow fractionation technique (AF4). The two different methodologies used in EV isolation yielded similar data in EV characterization.

### EV isolation using centrifugation

Cell culture media from HEK293T cells or J774.1 cells grown to 80% confluence in T75 flasks was harvested and undergone through multiple stages of centrifugation (Fig.1A).

EVs were isolated from the initial culture media samples using a series of centrifugation and ultracentrifugation steps. First, the samples were centrifuged at 2000g for 10 minutes to remove cell debris. The supernatant was then subjected to a centrifugation at 10,000g for 30 minutes (Rotor JA-25.50, Beckman counter, USA) to eliminate larger vesicles and other contaminants. Following this, the supernatant was ultracentrifuged at 110,000g for 120 minutes (Rotor SW32 Ti, Beckman counter, USA) to pellet the EVs. The resulting pellet was resuspended in phosphate-buffered saline (PBS) to a volume of approximately 25-30 ml. The resuspended pellet was then loaded onto a 30% sucrose cushion (4-5 ml per sample) and ultracentrifuged again at 110,000g for 120 minutes. After centrifugation, the upper fraction was removed, and the sucrose fraction was carefully collected. To remove sucrose, the fraction was diluted 10 times with PBS and subjected to another round of ultracentrifugation at 110,000g for 120 minutes. The final EV pellet was resuspended in PBS, with the volume adjusted to approximately 0.5 ml per sample. The concentration and size distribution of the isolated EVs were then measured using Dynamic Light Scattering (DLS).

### EV isolation using AF4

Cell culture media from HEK293T cells or J774.1 cells grown in CELLine Bioreactors was harvested once a week, cell debris were pelleted with centrifugation at 300g for 15 min., the supernatant containing EVs was concentrated using 100 kDa MWCO Amicon spin columns at 3000 g, 30min. Concentrated EV containing media was filtered with 0.45 µm syringe filter before EVs isolation with the technique of asymmetric flow field-flow fractionation (AF4, Wyatt Eclipse with Dilution Control Module, Wyatt Technology, Santa Barbara, CA). With AF4 separation we were able to concentrate, quantitate, characterize, and elute EVs, separate from contaminants, in real time with virtually no retention of contaminating soluble protein. Multiple detector readouts including Multi-Angle Light Scattering, Dynamic Light Scattering (MALS), and UV absorption at 280 nm used to monitor EVs and free proteins separation and EV size distribution in real time for downstream fractions collection. Different EV size fractions diluted with AF4 running buffer PBS were pooled together and concentrated using 100 kDa MWCO Amicon spin columns at 3000g, 30min. For downstream applications isolated EVs were kept in 0.05 µm filtered either in PBS or in HEPES-buffered artificial cerebrospinal fluid (ACSF) solution containing (in mM): 125 NaCl, 2.5 KCl, 10 HEPES, 1.5 MgCl_2_, 2.5 CaCl_2_, 10.0 glucose, and 10 Sucrose to adjust osmolarity to 290 mOsm, pH 7.3.

### EV characterizations

EV sizes were measured using dynamic light scattering (DLS) and nanoparticle tracking analysis (NTA) techniques. DLS EV size measurements were performed on vesicular fractions using the DynaPro NanoStar II (Wyatt Technology Corp., Santa Barbara, CA) in quartz cuvettes. The DLS data was analyzed using Dynamics Touch software (Wyatt Technology Corp). NTA measurements were performed at the University of Chicago Biophysics core using NanoSight NS300 (Malvern Panalytical Ltd, UK).

### Zeta potential

Nanoparticles zeta potentials were measured using DynaPro ZetaStar ( Waters I Wyatt Technology, Santa Barbara, CA USA).

### Cryo-EM sample preparation and data acquisition

Cryo-electron microscopy (Cryo-EM) samples of both EVs and liposomes were prepared using Lacey carbon EM grids (200 mesh) (Electron Microscopy Sciences, USA). The grids were first glow-discharged for 30 seconds at 20 mA in a Pelco EasiGlow system. Subsequently, 3.5 µL of the sample’s aqueous solution was applied to the carbon side of the EM grids. The grids were incubated in the instrument’s humidity chamber at 100% humidity and room temperature for 3 minutes. After incubation, the grids were blotted for 3.0 seconds at a blot force of -1 and then plunge-frozen into precooled liquid ethane using a Vitrobot Mark IV (FEI, USA).

Cryo-electron micrographs of vitrified samples were acquired using a Glacios Cryo-TEM transmission electron microscope (Thermo Fisher Scientific, USA) equipped with a STEM detector and operating at an accelerating voltage of 200 kV. Grid mapping and image acquisition were conducted using EPU Software, with nominal magnifications set to 180× for grid mapping and 13,500× for image acquisition. High-magnification images were captured at a nominal magnification of 73,000×, corresponding to a pixel size of 0.2 nm, with a defocus value of −5.5 µm.

### Immunogold Staining of Vesicles for Cryo Microscopy

EVs and HEVs were subjected to immunogold labeling for visualization under Cryo Microscopy. For immunostaining, the vesicle samples were incubated with primary anti-CD63 antibodies (Thermo Fisher) at a dilution of 1:10 for 1 hour at room temperature (RT) while continuously agitated. Following incubation, excess primary antibodies were removed by ultracentrifugation at 150,000 g for 45 minutes at 4°C, using a Sorvall mX150 Micro-Ultracentrifuge (Thermo Fisher, USA) with Rotor S140AT (Thermo Fisher). The supernatant was discarded, and the pellet was resuspended in 50 µL of PBS buffer. Subsequently, secondary antibodies conjugated with 10 nm gold nanoparticles were added to the samples at a dilution of 1:50. The vesicles were incubated with the secondary antibody for an additional 1 hour at RT. The immunogold-labeled vesicles were then applied to plasma-cleaned 300 mesh carbon-coated grids (Electron Microscope Sciences, USA) and frozen using a Vitrobot system (Thermo Fisher, USA).The prepared samples were examined using the Thermo Scientific Glacios Cryo-TEM 200 kV, equipped with a cryo-autoloader.

### EV fluorescent labeling

A number of fluorescent probes were used to label EV membranes including: Bodipy-TR, Bodipy-FL, NBD, R18 and Abberior STAR RED^TM^ membrane dyes. In most cases, the dyes incubation with EVs for 30 minutes was followed by unincorporated dyes removal using either Exosomes Spin Columns MW 3000 (Invitrogen, Thermo Fisher Scientific, USA) or Zebra^TM^ Spin Desalting Columns 7K MWCO (Thermo Fisher Scientific, USA).

### Liposomes and DOTAP LNP Production

For the production of LNPs with a single lipid composition, we employed a standard liposome preparation method. Lipids dissolved in chloroform were dispensed into a round-bottom flask using solvent-safe tips. To prevent oxidation, the remaining stock solution was purged with argon. The flask was then connected to a Rotavapor R-200 (Büchi, Germany) rotary evaporator and subjected to evaporation under vacuum at room temperature with slow rotation for 1 hour. Following solvent evaporation, the flask containing the lipids was kept under vacuum overnight to ensure complete drying.

A buffer of choice, either PBS or ACSF, was added to the flask and the mixture was sonicated for 30 minutes at 40°C (37 kHz frequency, 100% power). For mRNA loading experiments, mRNA was added to the sonicated mixture. The resulting preparation was extruded 20 times through a polycarbonate membrane (0.1–0.2 µm pore size) using an Avanti Mini Extruder (Avanti Research, Alabaster, Alabama, USA) to achieve uniform liposome formation and efficient RNA encapsulation.

### Complex LNP Production

Unless otherwise stated, Avanti’s LNP Lipid Blend 2 mixture (Avanti cat. number: 300213), composed of the following components: 49 mol% DOTAP, 38.5 mol% Sitosterol, 11 mol% DSPC, and 1.5 mol% DMG-PEG2000 - dissolved in ethanol, was used for LNP production. The mRNA of choice, dissolved in 10 mM citrate buffer prepared with RNase-free water, was combined with the lipid mixture. Calculations for the desired N/P ratio were performed, and the calculated volumes of the Lipid Blend 2 mixture and mRNA solution were mixed in a single-use microfluidic chamber of the NanoAssemblr™ Ignite™ nanoparticle formulation system (Cytiva, USA) following the manufacturer’s suggested injection protocol.

Subsequently, ethanol and citrate buffer were removed from the resulting LNP mixture and replaced with a buffer of choice using a 100 kDa MWCO Amicon Ultra Centrifugal Filter (Millipore, USA).

On a few occasions, LNPs were prepared using a homebuilt Jets Mixer for Flash NanoPrecipitation, as described in the literature ^22^.

### Production of HEVs by extrusion

Lipids were initially dissolved in chloroform and transferred to a round bottom flask. The organic solvent was then evaporated at room temperature using a rotary evaporator (RotoVap, Büchi Labortechnik, Flawil, Switzerland) under vacuum, resulting in the formation of a lipid film. This lipid film was subsequently dissolved in artificial cerebrospinal fluid (ACSF). Following hydration, the lipid suspension was sonicated at 40°C for 50 minutes at a frequency of 37 kHz and a power setting of 100. The sonicated suspension was then filtered by centrifugation at 300g for 5 minutes using a 0.45 µm centrifuge filter tube. The lipids were further processed by extrusion through a 0.1 µm polycarbonate filter for 20 cycles using an Avanti Hand Extruder (Avanti Polar Lipids). The concentration of the resulting liposomes was measured, and the sample was then mixed with an extracellular vesicle (EV) sample at a specified ratio. This mixture was subjected to a second extrusion through a 0.2 µm polycarbonate filter. Finally, the concentration and polydispersity index (PDI) of the obtained vesicles were analyzed using DLS.

### *In vivo* toxicity of EVs, Liposomes, LNPs, and HEVs

Adult (3 to 6 months old) C57BL/6 mice were anesthetized using Ketamine/Xylazine anesthesia and 100 µl of solution containing combination of nanoparticles and 100 µM of Propidium Iodide was instilled intratracheally to the left lung lobe using bended plastic catheter. After 1 hour mice were sacrificed by combination of the anesthetic overdose and cervical dislocation followed by intratracheal injection of 1.5 ml of low gelling temperature agarose (SeaPlaque^TM^ GTG^TM^ Agarose, Lonza, USA) dissolved in 4% PFA in PBS to keep lung volume. The mouse body was cooled down by placing in ice, than lung was dissected out of the body and placed overnight in 4% PFA at +4°C. Next day 200µm thick lung slices were cut using vibratome Leica VT 1000S (Leica, Germany). The lung slices were incubated for 30 min with NucBlue^TM^ Live Reagent^TM^ (Hoechst 33342) (ThermoFisher Scientific, USA) to label cells nucleus followed by laser confocal imaging using Leica SP5 inverted microscope (Leica, Germany).

### Microscopy

Laser confocal microscopy and super resolution STED imaging were performed in the Integrated Light Microscopy Core at University of Chicago, using a Leica SP5 and Leica SP8 (Leica, Germany) inverted laser confocal microscopes with STED functionality.

### Super-resolution STED EV imaging and analysis

For super resolution imaging EVs were labelled with Abberior STAR RED^TM^ membrane dye (Abberior, Gottingen, Germany) and 100nm FluoSpheres^TM^ carboxylate, yellow-green (ThermoFisher Scientific, USA) were encapsulated in DOTAP liposomes. The nanoparticles were imbedded in ProLong^TM^ Diamond Antifade mountant (ThermoFisher Scientific, USA) at least 24 hours before imaging. STED depletion was performed with 775 nm laser set at 13.8% power. STED images were analyzed using the Leica LIGHTHING (Lng) image deconvolution technique and further analyzed using ImageJ analysis software.

### SDS-Polyacrylamide Gel Electrophoresis (SDS-PAGE) and Western Blot Analysis

EV were mixed with 5X Laemmli buffer (250 mM Tris-HCl, pH 6.8, 10% SDS, 30% glycerol). Addition of DTT as reducing agent in buffer and heating at 95°C for 5 minutes depended on the analyzed proteins. Samples were then separated using Mini-protean TGX precast 4-20% gradient polyacrylamide gels. Electrophoresis was performed in Tris-Glycine running buffer at a constant current of 25 mA for 60 minutes.

The resolved proteins were transferred onto 0.45 µm nitrocellulose membranes using a Trans Blot SD semi-dry transfer cell (Bio Rad) at a constant voltage of 15 V for 30 minutes. Membranes were blocked for 1 hour at room temperature in 1X TBST (containing 0.1% Tween-20) with 5% (w/v) skim milk.

Primary antibody incubation was carried out overnight at 4°C using the following antibodies diluted in blocking solution: CD63 (H5C6, BD Biosciences, #556019, 1:500), CD81 (M38, ThermoFisher, cat. # 10630D, 1:1000), TSG101 (EPR7130(B), Abcam, #ab125011, 1:1000).

After incubation, membranes were washed three times for 5 minutes each with TBST solution. Secondary antibody incubation was performed for 1 hour at room temperature using HRP-conjugated goat anti-rabbit IgG and HRP-conjugated goat anti-mouse IgG (Abcam) 1:5000 + 1:20000 StrepTactin-HRP Conjugate (Bio Rad) diluted in 5% blocking solution.

Blots were washed three times in TBST solution. Protein bands were visualized using the c600 Imaging System (Azure Biosystem) for chemiluminescence.

### Flow cytometry

Cell viability was analyzed by flow cytometry. Briefly, HEK293T cells were seeded in 24-well plates at a cell density of 5x10^4^ and 24 hours after seeding, were incubated for three hours following exposure to increasing amounts of either EVs, LNPs, Liposomes or HEVs, as stated in the associated figure legend. At the end of the incubation period, particles were removed, and cells were gently detached by treatment with Accutase^TM^ and transferred into flow tubes. Cells were gently washed twice with cold PBS without calcium and magnesium supplemented with 0.2 % of BSA. Subsequently, cells were resuspended in 200 µl of PBS without calcium and magnesium and 0.2% BSA and stained with propidium iodide (PI, 1 µM) immediately prior to analysis. Cells were gated based on their forward and scatter properties (FSC and SSC, respectively) to exclude cell debris and doublets. Toxicity was evaluated as the percentage of PI+ cells and fusion efficiency was determined by gating NBD+ cells within the PI-cell population. Images were acquired at the University of Chicago Cytometry and Antibody Technology core facility with the Novocyte platform using Agilent Penteon 5-30 and processed by analysis with FlowJo Software.

### FRET measurements

Kinetics FRET measurements were performed at the University of Chicago Biophysics core facility using modular spectrofluorometer HORIBA Fluorolog-3 (HORIBA, Kyoto Japan). Steady state FRET measurements were performed using BioTek Synergy Mx multi-mode microplate reader ( BioTek Instruments Inc., Winooski USA).

### Statistics and Reproducibility

To analyze the statistically significant difference between the groups, ANOVA analyses (Student t-test and Mann-Whitney Rank Sum Test) were used. Data are represented as mean ± SEM. At least 3 mice were used in all experimental studies.

## Data Availability

The authors declare that the data supporting the findings of this study are available within the paper and supplementary information files. All source data underlying the graphs shown in the main text.

## Acknowledgements

We thank The University of Chicago core facilities: Integrated Light Microscopy Core, Cytometry and Antibody Technology core, Biophysics core facility, Advanced Electron Microscopy Core Facility for technical help and access to instruments.

The authors thank Prof. Stephen C. Meredith (The University of Chicago) for help with LNP preparations, Dr. Adam Brooks and Waters I Wyatt Technology Corporation for kind help with Zeta potential measurements, Aniruddhsingh Solanki (The University of Chicago) for technical help with mouse in-vivo injections, Abberior Instruments America for providing free samples of Abberior STAR fluorescent dyes and Prof. Paul Schlesinger (Washington University in St. Louis) for helpful discussions and advice on liposome preparation.

This work was supported by NIH/NHLBI grant R01HL125076 to DJN.

## Author contributions

Deborah J. Nelson, Conceptualization, Supervision, Funding acquisition, Investigation, Manuscript Writing and editing; Vladimir Riazanski, Investigation, Data acquisition and analysis, Manuscript Writing; Lada Purvina, Data acquisition and analysis, Drafting article; Luca Cavinato, Data acquisition and analysis; Zihao Sui, Data acquisition and analysis; Lauria Sun, Data acquisition and analysis.

